# Introgression impacts the evolution of bacteria, but species borders are rarely fuzzy

**DOI:** 10.1101/2024.05.09.593304

**Authors:** Awa Diop, Louis-Marie Bobay

## Abstract

Most bacteria engage in gene flow and that this may act as a force maintaining species cohesiveness like it does in sexual organisms. However, introgression (gene flow between the genomic backbone of distinct species) has been reported in bacteria and is associated with fuzzy species borders in some lineages, but its prevalence and impact on the delimitation of bacterial species has not been systematically characterized. Here, we quantified the patterns of introgression across 50 major bacterial lineages. Our results reveal that bacteria present various levels of introgression, with an average of 2% of introgressed core genes and up to 12% in *Campylobacter*. Furthermore, our results show that some species are more prone to introgression than others within the same genus and introgression is most frequent between highly related species. We found evidence that the various levels of introgression across lineages are likely related to ecological proximity between species. Introgression can occasionally lead to fuzzy species borders, although many of these cases are likely instances of ongoing speciation. Overall, our results indicate that introgression has substantially shaped the evolution and the diversification of bacteria, but this process does not substantially blur species borders.

## Main

Species borders are important to identify and yet difficult to define ^1–3^. Because bacteria are asexual organisms, they do not easily adhere to the species concepts developed for sexual organisms, i.e., the Biological Species Concept (BSC) and related definitions ^4,5^. However, a growing body of evidence is supporting the idea that the vast majority of bacteria do engage in gene flow and that the BSC may be applicable to these organisms ^6–12^. Through diverse routes and vessels, bacterial cells are capable of exchanging genetic material and the transferred DNA may be related enough to sequences of the recipient genome to be swapped through homologous recombination. On the other hand, other transfer events—termed Horizontal Gene Transfers (HGTs)—do not require sequence relatedness or can rely on microhomologies for DNA integration ^13–15^. From an evolutionary point of view, homologous recombination promotes the exchange of alleles between highly related homologous sequences, and this process is comparable to gene flow in sexual organisms. In contrast, HGT can lead to the gain of new genes previously absent in the recipient genome and can occur between more distantly related—or unrelated—sequences ^13–15^. For the purpose of species definition, we and others have focused primarily on the patterns of gene flow (i.e., homologous recombination) as a force maintaining the genetic cohesiveness of bacterial species ^8,10,12,16,17^. However, gene flow can also have the opposite effect: blurring species boundaries.

Earlier studies relying on multi-locus sequence typing (MLST) first observed clear discrepancies across sequence markers while attempting to classify and type bacterial strains of several lineages ^18–23^. Notoriously, strains of the genus *Neisseria* were found to form “fuzzy” species due to the recombinogenic nature of this lineage ^22,24,25^. Additional cases of species fuzziness were reported in subsequent MLST studies ^19,20,23^, but the advent of full genome sequencing solved the issue of incongruent gene marker phylogenies by building phylogenetic consensuses across hundreds or thousands of genes ^26–28^. Nevertheless, these studies provided some of the first evidence that gene flow can be somewhat porous across bacteria and that bacterial species borders may be fuzzy in some lineages ^29,30^.

Gene flow is pervasive between closely related genomes of most species and rarely occurs between genomes showing more than 2 to 10% nucleotide divergence ^12^. This restriction is thought to reflect the mechanistic constraints of the homologous recombination machinery, which requires the presence of stretches of identical nucleotides shared between the recipient strand and the incoming strand ^31^. However, gene flow can occasionally occur between more distantly related genomes ^12,22,24,32,33^. Such patterns were clearly detailed in a study comparing *Campylobacter coli* and *Campylobacter jejuni*, where a substantial portion (∼20%) of the genome of these two species appears to be engaging in gene flow, while the rest of the genome is substantially diverged (∼85% sequence identity) ^33^. Importantly, these exchanges of DNA occur through allelic replacements in the genomic backbone of these species (i.e., the core genome) and do not occur through the transfers and gains of accessory genes *via* HGT which are frequently observed between distant lineages. Such patterns of gene flow between distinct species are reminiscent of introgression in sexual organisms. This process is likely responsible for the species fuzziness observed in some bacterial lineages, but the prevalence and the impact of introgression on the delimitation of bacterial species has not been systematically assessed.

Here, we characterized the patterns of introgression across 50 major bacterial lineages. Our results indicate that introgression is common in core genomes across bacteria. However, introgression tends to be highly overestimated in several species due to the inaccuracy of species borders’ definition. Among the analyzed lineages, the genera *Campylobacter*, *Haemophilus* and *Escherichia-Shigella* presented the highest levels of introgression. Overall, our analysis indicates that bacterial species present various levels of fuzziness across lineages and these could correspond to ongoing speciation events. However, most species appear clearly delineated based on core genome phylogenies.

## Results

### Prevalence of introgression across bacterial lineages

We first quantified the amount of DNA exchanged by gene flow between the core genome of distinct species (i.e., introgression). We built and analyzed the core genome of 50 bacterial genera and classified all genomes within each genus into ANI-species based on the pairwise ANI of core genomes using a cutoff of 94% sequence identity as performed previously ^34^. We generated a maximum-likelihood phylogenomic tree for each genus using the concatenated core genome alignments. The trees segregated the vast majority of the ANI-species into monophyletic groups (i.e., phylogenetic species) (Table 1 and Supplementary Fig. 1). To make a distinction between gene flow *within* species and gene flow *between* species, we refer to gene flow between species as *introgression* to parallel the processes described in sexual organisms. To assess the prevalence of the patterns of introgression between ANI-species within each of the 50 genera included in this study, we used an approach based on phylogeny and sequence relatedness to detect and quantify introgression (see Methods). Briefly, for a given genus, we inferred introgression events based on the phylogenetic incongruency between gene trees and the core genome tree in a particular ANI-species. A gene sequence was inferred as introgressed between two ANI-species when forming a monophyletic clade that was inconsistent with the unrooted core genome phylogeny (Fig. 1). To be categorized as introgressed, a core gene sequence was also required to be statistically more similar to the sequence of a different ANI-species than its own (see Methods). For each ANI-species, levels of introgression were expressed as the fraction of core genes that satisfied these two criteria (Fig. 1).

**Fig. 1.**
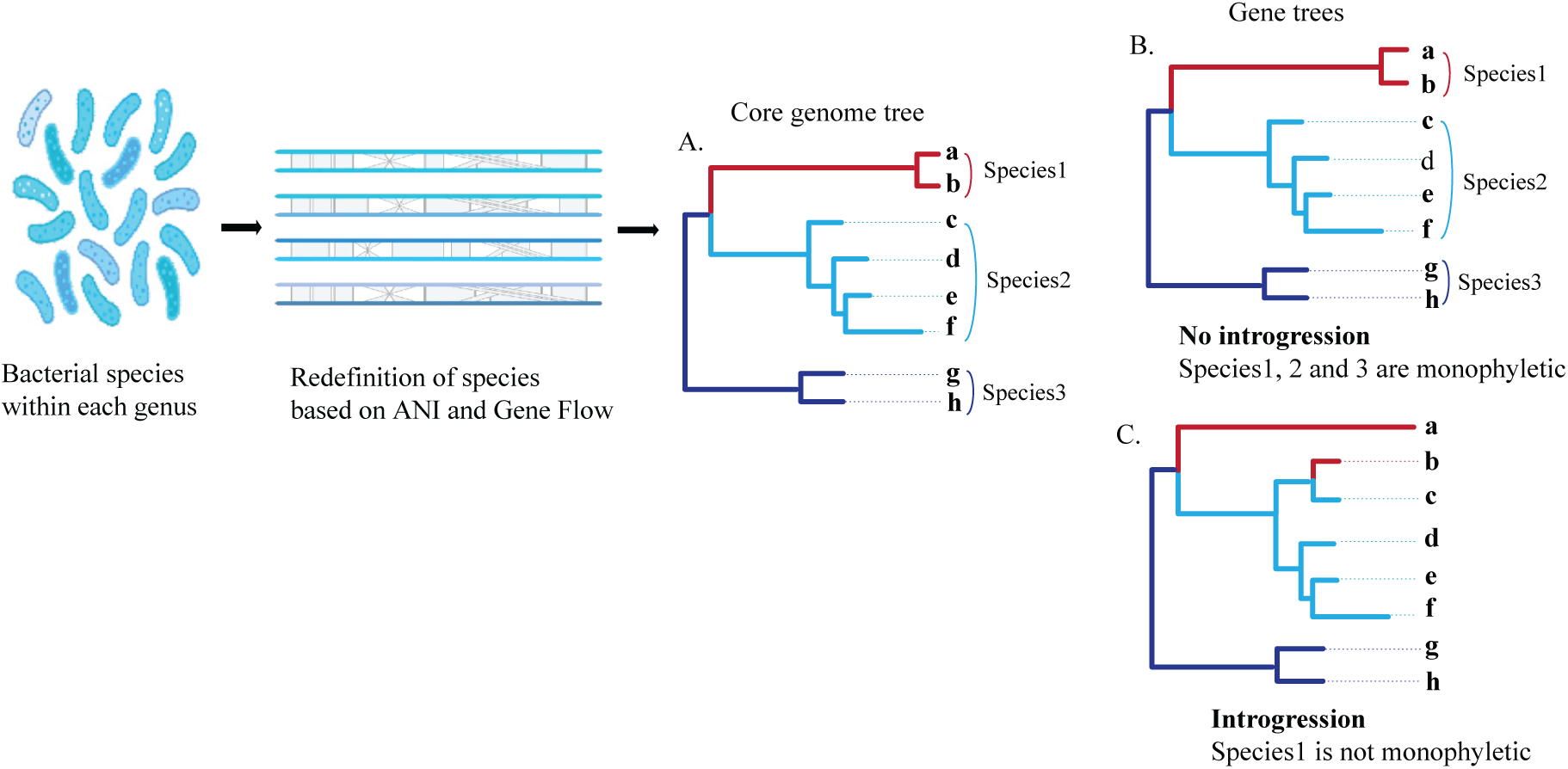
Approach used to infer introgression across lineages. Genomes within the same genus were considered part of the same species (ANI-species) when sharing at least 94% the Average Nucleotide Identity (ANI) across the core genome. Phylogenetic trees were inferred from the concatenated core genome alignment of each genus. **b**, and **c**, Phylogenetic trees were inferred for each core gene of each genus. Introgression events were inferred based on the phylogenetic incongruency between gene trees and the core genome tree in a particular ANI-species. On the example represented in this figure: the topology of tree ‘A’ is congruent with that of tree ‘B’ in all ANI-species. In contrast, the topology of unrooted tree ‘C’ is incongruent with the topology of the core genome tree ‘A’ for ANI-species 1 and 2. This case represents a potential introgression event from ANI-species 1 to ANI-species 2. Putative introgression events detected based on tree incongruencies were then inferred as introgression events if the relatedness of the introgressed sequence was statistically higher than the relatedness of the core genome of the two species (>2 S.D., see Methods).

Using this approach, we observed that the genera studied here present various levels of introgression with an average of 7% of introgressed genes across genera, suggesting that this process is rather common in bacteria (Fig. 2; Table 1; Supplementary Table 1).

**Fig. 2.**
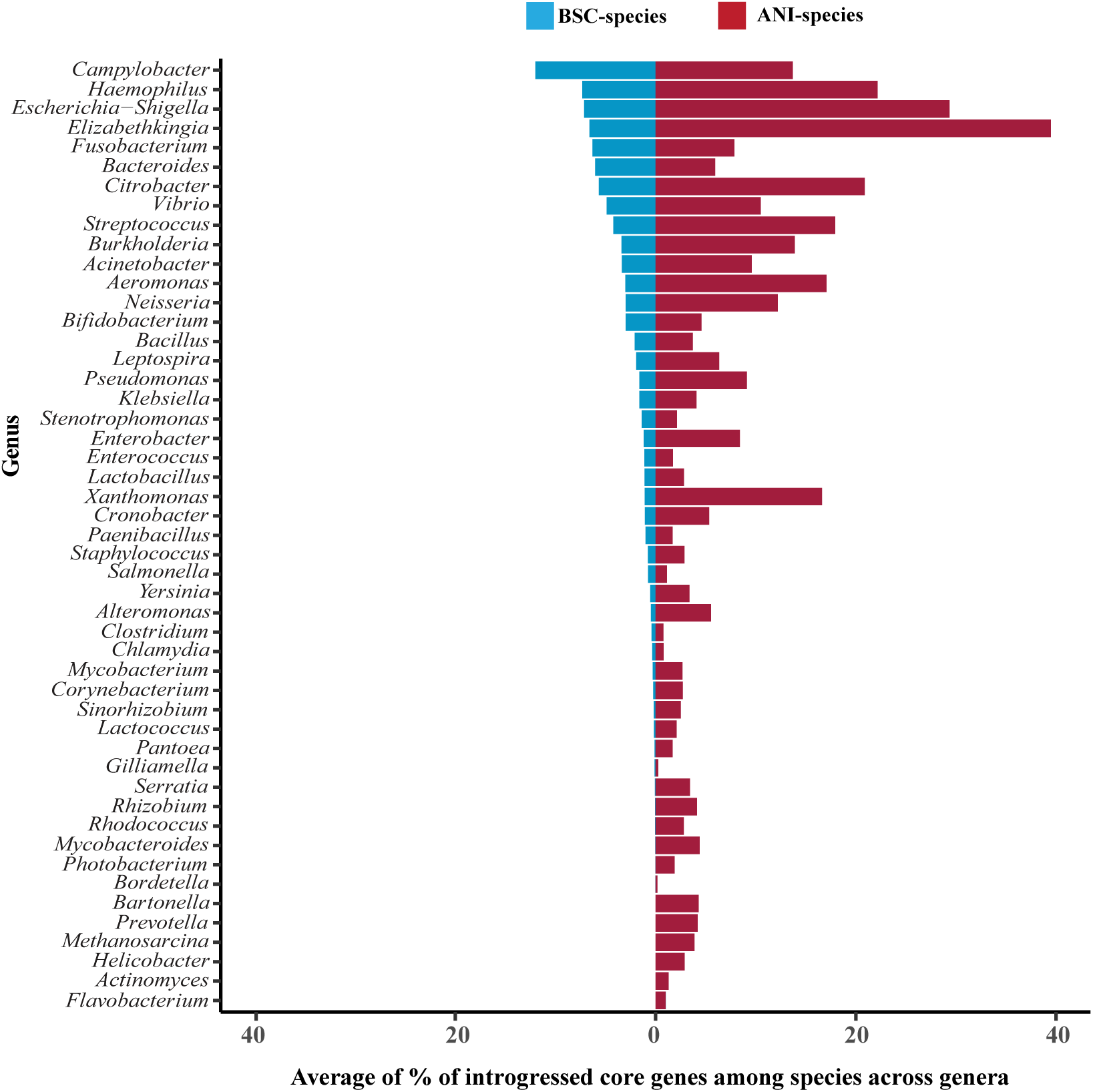
Levels of introgression across genera. Levels of introgression were expressed as the average percentage of introgressed core genes between all pairs of species within each genus. The average percentage of introgressed core genes of each genus were estimated by classifying species into ANI-species (red) or by classifying species into BSC-species (blue). ANI-species were defined using a cut-off of ≥94% along the core genome. BSC-species were defined as groups of genomes showing significant levels of gene flow based on the signal of homoplasic alleles (see Methods).

The previous results relied on the ANI as an empirical approach to define species. However, ANI-based methods are not anchored in a solid theoretical framework and we have previously suggested that gene flow could be used to refine ANI-species borders following similar principles as in the Biological Species Concept (i.e., BSC-species)^12^. Therefore, the levels of introgression estimated above may be inflated by the inaccurate identification of species borders (i.e., if two ANI-species may in fact represent a single BSC-species). Indeed, our results indicate that most of the introgression events occurred between closely related or sister ANI-species based on both the core genome phylogenies and the relatedness of the core genomes (Supplementary Fig. 1, Figs. 2a and 2b and Table 2) and it is therefore very likely that slightly adjusting species borders would reveal different estimates of introgression levels. To address this issue, we refined the borders of ANI-species based on the patterns of gene flow to generate BSC-species based on the signal of homoplasic alleles relative to non-homoplasic alleles (*h/m*), as previously described ^12^ (and see Methods). Following this approach, we found that most of the closely related ANI-species that shared high levels of introgression were in fact classified into a single BSC-species (Supplementary Tables 2 and 3). In most cases, BSC-species would match ANI-species borders by slightly adjusting the ANI threshold used to define species borders (note that, however, we cannot predict how the ANI threshold needs to be adjusted as it appears to be lineage- or species-specific). For instance, *Streptococcus parasanguinis* ANI-sp19 presented 37.8% of its core genome introgressed with *Streptococcus parasanguinis* ANI-sp47, but our analyses revealed that these two species form in fact a single BSC-species based on the signal of gene flow (these ANI-species were slightly under the 94% ANI threshold used to define ANI-species). Another example is the genus *Elizabethkingia* in which some ANI-species shared some of the highest levels of introgression, but our analyses indicate that these ANI-species are part of the same BSC-species (BSC-sp3) (Supplementary Fig. 1r and Tables 1, 2 and 3). ANI-sp3 (composed of *E. miricola* and *E. bruuniana*) presents ∼19% of its core genome introgressed with ANI-sp1 (composed of *E. occulta* and *E. ursingii*), but these ANI-species were highly related, sharing between 92 – 93% sequence identity). These results indicate that, in many cases, the higher levels of introgression detected between ANI-species do not correspond to true introgression events once we redefined species based on the signal of gene flow but rather correspond to frequent homologous recombination events among genomes within the same (biological) species.

After redefining ANI-species into BSC-species (Supplementary Table 2), we focused our analysis on the pairs of BSC-species where both BSC-species are composed of at least 15 genomes, to further ascertain that these species are truly distinct (low genome numbers affect our ability to reclassify species based on gene flow). Our results revealed that introgression impacts on average 2% of the core genome of bacterial species. Across genera, introgression ranged from 0% in six genera to 12% in *Campylobacter (*Fig. 2 and Table 1 and Supplementary Table 1). Within each genus, BSC-species displayed various levels of introgression, where 0 to 59% of core genes were found to be introgressed between BSC-species (Fig. 3a and Supplementary Table 1). These results indicate that a few bacterial species are very fuzzy even after redefining species based on gene flow. Among the analyzed lineages, *Campylobacter*, *Haemophilus* and *Escherichia-Shigella* present the highest levels of introgression (Fig. 2 and Table 1 and Supplementary Table 1) with 12, 7.3 and 7.1% of their core genome introgressed, respectively.

**Fig. 3.**
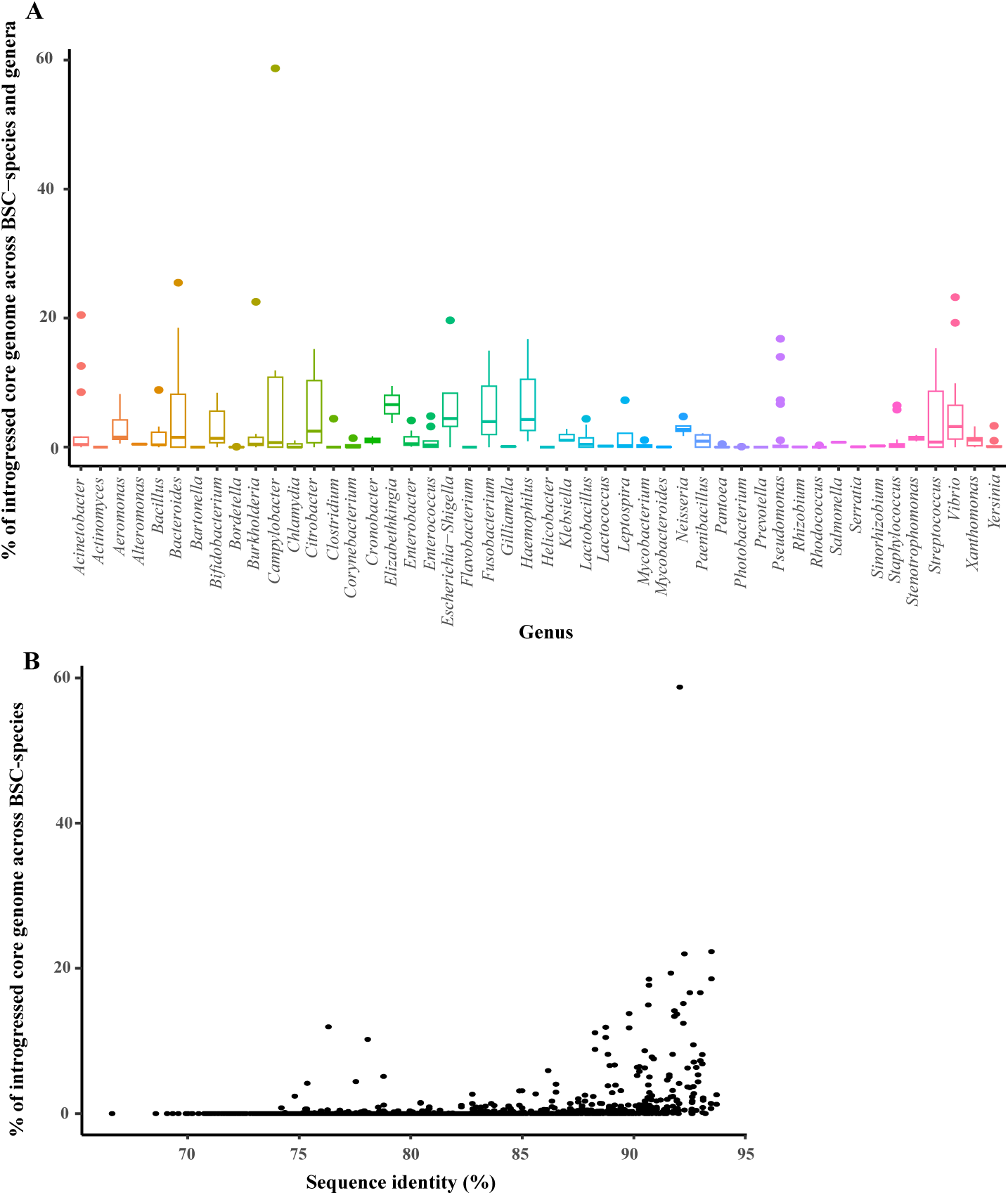
Levels of introgression across BSC-species and genera. **a**, Percentage of introgressed core genes across pairs of BSC-species within each genus. **b**, Relationship between levels of introgression and core genome sequence identity across pairs of BSC-species.

**Fig. 4.**
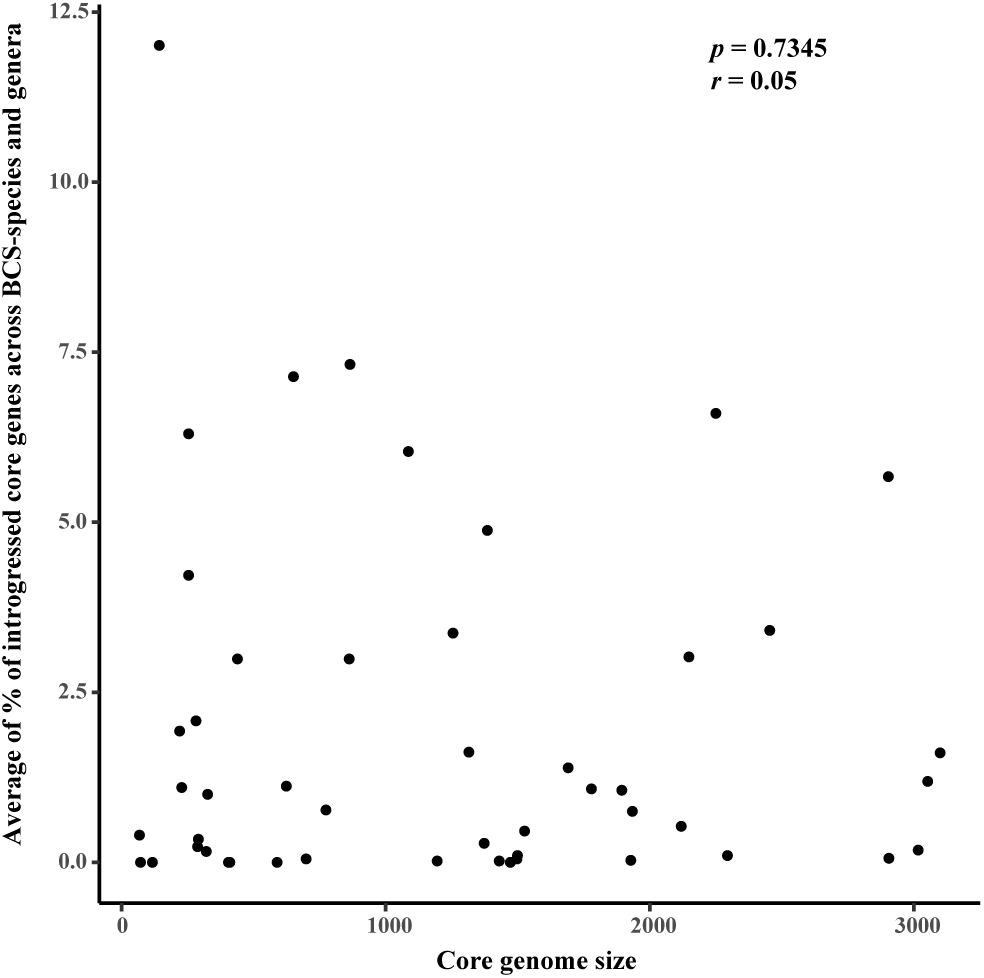
Correlation between core genome size and levels of introgression across genera. The levels of introgression were represented by the average percentage of introgressed core genes within each genus. Correlation was assessed using Spearman’s correlation coefficient *Rho* and associated *p*-value (*P*).

The distribution of the levels of introgression across genera showed that most of the BSC-species within the same genus share between 0 and 6% of introgressed core genes (Supplementary Fig. 3). These results indicate that substantial amounts of introgression are commonly occurring between related bacterial species. However, levels of introgression can vary sharply across species of the same genus (Fig. 3a and Supplementary Table 1). An illustrative example is the genus *Campylobacter* where species share between 0 to 59% of introgressed core genes. Similar trends were observed in several genera such as *Bacteroides, Vibrio*, *Burkholderia, Acinetobacter*, *Escherichia-Shigella*, *Pseudomonas*, *Haemophilus,* and *Streptococcus*, where species share typically between 0 to 20% of introgressed core genes, respectively (Fig. 3a and Supplementary Table 1).

### Introgression is common between species sharing the same ecological niche

Our results suggest that some species are more prone to introgression than others within the same genus. This may be related to the ecological and physical proximity of these species which would increase opportunities for genetic exchange between them. This is the case of the genus *Haemophilus* whose species are predominantly human symbionts ^35–38^. For instance, BSC-sp2 (composed of *H. influenzae,* and *H. aegyptius*) presents 17% of its core genome introgressed, among which 16.7% are shared with its most closely related species *H. haemolyticus* (BSC-sp1), both species co-colonizing mainly the human respiratory tract (Supplementary Fig. 1x, Tables 1 and 2). This is also the case for the group *Escherichia-Shigella* which includes species that colonize the gut of a broad range of animals including the human gut ^39–41^. It has been shown that recombination and introgression are important forces shaping the genomic evolution and diversification of *E. coli* ^30,39^*. E. fergusonii* (BSC-sp4) presents 20% of its core genome introgressed, most of which is shared with BSC-sp2 (composed of *E. coli, S. sonnei, S. dysenteriae, S. boydii and S. flexneri*) (Supplementary Fig. 1q, Tables 1 and 2), its close relative that also colonizes animal guts ^39,41–44^. The genus *Campylobacter* constitutes another example with *C. coli* (BSC-sp28) and *C. jejuni* (BSC-sp3) colonizing the same body site (mainly the gastrointestinal tract) in humans and livestock (Supplementary Fig. 1k and Table 2). Consistent with previous studies ^32,33,45^, our results showed high levels of introgression between *C. coli* (BSC-sp28) and *C. jejuni* (BSC-sp3): 11% of core genes of *C. jejuni* have been introgressed with *C. coli* and 12% of core genes of *C. coli* have been introgressed with *C. jejuni* (Table S2). In addition to *C. coli* and *C. jejuni*, the genomes of *C. concisus* are classified into two distinct biological species (BSC-sp 24 and BSC-sp4, respectively) and present introgression of 59% of the core genome.

We observed that higher levels of introgression are typically occurring between closely related BSC-species (Main text Fig. 3b and Supplementary Fig. 1 and Table 2) and we have previously hypothesized that the ability to engage in gene flow is linked to the ability of the recombination machinery to initiate homologous recombination more efficiently when sequences share high identity^12^. Most of the species within the genus *Streptococcus* that share the highest levels of introgression are highly related to one another. For instance, BSC-sp2 (composed of *S. lutetiensis* and *S. equinus*) presents 14% of introgressed core genes shared with *S. equinus* (BSC-sp70) (these species shared between 91 and 93% ANI). *S. pyogenes* (BSC-sp45) and *S. dysgalactiae* (BSC-sp7) which are close relatives (shared between 89 to 91% ANI) constitute another example within this genus. Both species share over 12% of introgressed core genes with one another (Supplementary Fig. 1at, Tables 1 and 2). The same trend was observed in the genus *Bacteroides* where many species share higher levels of introgression with their close relatives. For instance, introgression between closely related clades BSC-sp1 (composed of *B. intestinalis*, *B. timonensis* and *B. cellulosilyticus*) and BSC-sp22 (*C. intestinalis*) was observed: 18% of the core genes of BSC-sp1 have been introgressed with *C. intestinalis* and 19% of the core genes of *C. intestinalis* have been introgressed with BSC-sp1 (Supplementary Fig. 1f and Table 2). In contrast the higher levels of introgression (10% of introgressed core genes) between BSC-sp23 (composed of *B. xylanisolvens, B. ovatus, B. caecimuris, B. koreensis* and *B kribbi*) and BSC-sp5 (*B. fragilis*), which are very distant relatives (both species shared between 78 to 84% ANI), constitute an exception within this genus. A similar exception was observed in the genus *Acinetobacter* where *A. baumannii* (BSC-sp38) shares higher levels of introgression (12%) with *A. radioresistens* (BSC-sp14), a distant relative (sharing only between 76 to 79% ANI) (Supplementary Fig. 1a and Table 2).

Within some lineages, most of the commensals and opportunistic pathogens present higher levels of introgression relative to pathogens, further suggesting that the frequency of introgression is dependent on the ecology of the species and is potentially adaptive. Species within the genus *Vibrio* are almost all free-living bacteria that often undergo variable selective pressures due to environmental changes, including seasonal changes but can become opportunistic pathogens in humans ^46^. Within this genus, BSC-sp59 (*V. diabolicus* and *V. antiquarius*), which is described as “Vibrio-related probiotic with no pathogenic antecedent” ^47,48^, presents 24% of introgressed core genes. In addition, the opportunistic human pathogens *V. cholerae* (BSC-sp63) and *V. alginolyticus* (BSC-sp20) ^46^ display higher levels of introgression with 10% and 19% of their core genome introgressed, respectively. In contrast, BSC-sp58 (composed of the human pathogens *V. vulnificus* and *V. fluvialis*) and BSC-sp32 (*V. parahaemolyticus*) present lower levels of introgression (1.6 and 1.9% of introgressed core genome, respectively) (Supplementary Fig. 1au and Table 1). A similar trend was observed within the genus *Acinetobacter* where *A. baumannii* (BSC-sp38) and *A. nosocomialis* (BSC-sp4) present the highest level of introgressed core genes within this genus (20% and 13% of introgressed core genes, respectively) (Supplementary Fig. 1a and Table 1). However, some genera are notable exceptions to this trend: for instance, the genus *Serratia*, which includes the opportunistic pathogen *Serratia marcescens* (BSC-sp2), has a lower incidence of introgression with 0.1% of its core genome introgressed.

### Introgression variation by gene function

Because introgression events are potentially adaptive, we analyzed the function of all introgressed genes. All core gene families were classified into COG (Categories of Orthologous Groups) categories using EggNog ^49^ (Supplementary Fig. 5). We compared the representation of COG categories in the introgressed genes relative to the entire core genome (Supplementary Fig. 6) among all genera (Fig. 5).

**Fig. 5.**
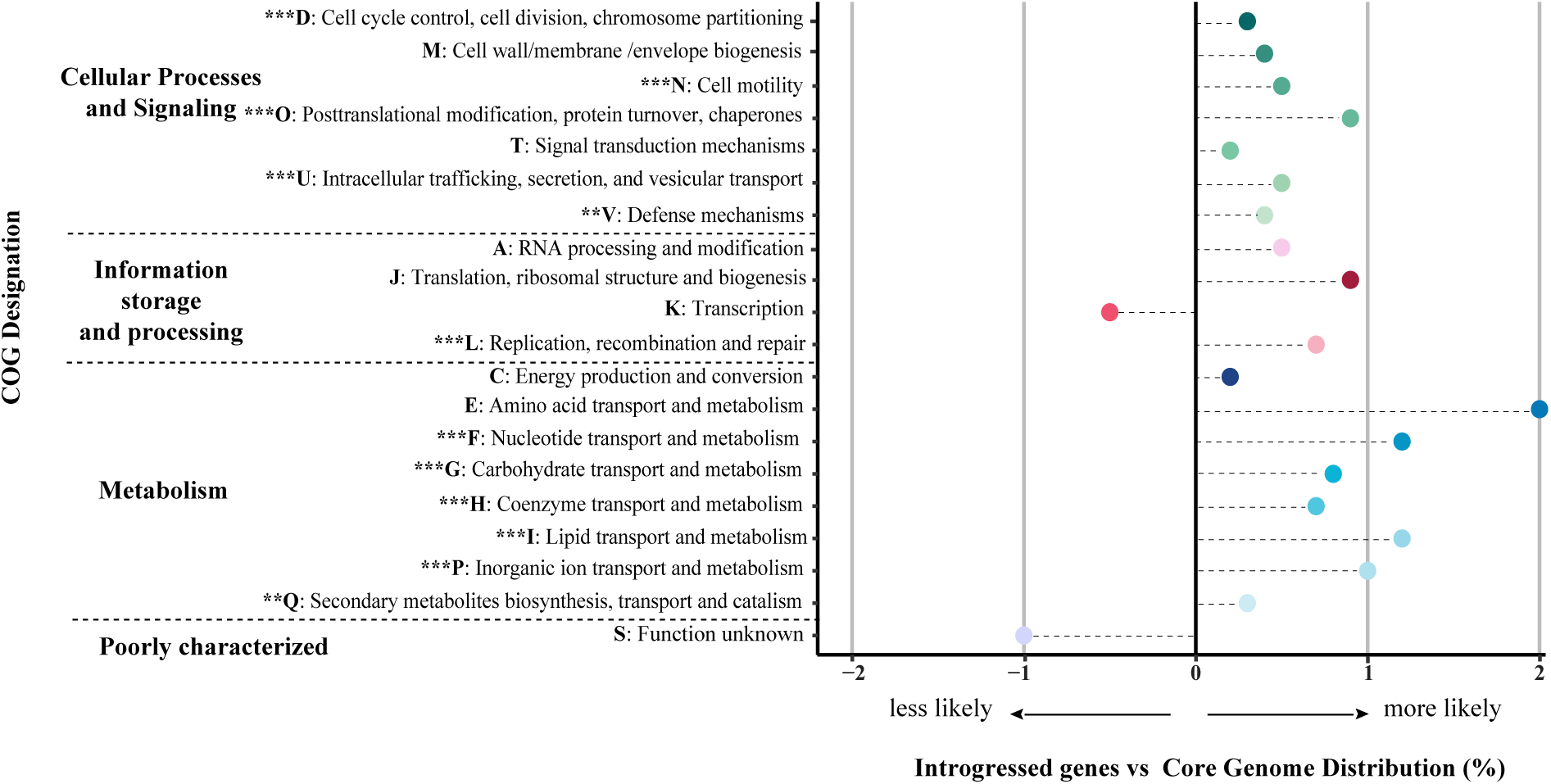
Functions of introgressed genes relative to the entire core genome across all genera. *P* values of COG categories are for the statistical test to detect a difference in COG categories using Wilcoxon test and were adjusted for multiple testing with a Bonferroni correction. See the legend key for the description of each COG category. COG Categories significantly less or more frequently introgressed relative to the core genome are noted as follow: * *P* < 0.05, ** *P* < 0.01 and *** *P* < 0.001.

Several functional categories were found to be introgressed at higher frequency relative to their representation in the core genome (Wilcoxon test with Bonferroni adjustment). Genes involved in carbohydrate transport and metabolism were found to be the most significantly overrepresented in introgression events (*P*<10^-7^). Other functional categories were found to be more introgressed than expected: Lipid transport and metabolism (*P*<10^-6^), coenzyme metabolism and transport (*P*<10^-5^), inorganic ion transport and metabolism (*P*<10^-5^), cell cycle control, cell division, and chromosome partitioning (*P*<10^-4^), intracellular trafficking, secretion and vesicular transport (*P*<10^-4^), cell motility (*P*<10^-3^), posttranslational modification, protein turnover, chaperones (*P*<10^-3^), nucleotide transport and metabolism (*P*<10^-3^), secondary metabolites biosynthesis, transport and catabolism (*P*<10^-2^), and defense mechanisms (*P*<10^-2^). Interestingly, in the “information storage and processing” category: Replication, recombination and repair was the only functional category found to be over-represented (*P*<10^-4^) (Fig. 5). The overrepresentation of functional categories in introgressed genes is presented in detail for each genus independently (Supplementary Fig. 6). The higher frequency of transfers of certain functions likely represents the role these introgression events play in bacterial adaptation. Indeed, introgressed genes are most frequently involved in carbohydrate and lipid metabolism, and recombination within these genes could be adaptive when colonizing a new environment or during fluctuations of resource availability.

## Discussion

Our results report, overall, that introgression is prevalent across bacterial lineages. All 50 genera that we studied showed evidence of introgression, although some lineages were much more prone to introgression than others. Our results support previous findings: introgression is common in *Neisseria* (although we observed lower levels of introgression because *N. gonorrhoeae* and *N. meningitidis* were classified into a single species) ^12,50^, *Campylobacter* ^12,32,33,45,51^*, Escherichia-Shigella* ^30^ and *Streptococcus* ^52,53^. Several species are indeed fuzzy, although the delimitation of species borders is possible, and defining species based on gene flow helps with these cases. For instance, frequent introgression within *Neisseria* makes it difficult to delimitate this genus into distinct species. However, inconsistent species delineation has been notoriously problematic in this genus: we classified *N. mucosa*, *N. sicca*, *N. subflava* and *N. perflava* into two different clusters (BSC-sp1 and BSC-sp5) and several closely related strains are currently assigned to different named species (Supplementary Fig. 1ag and Table 1). Additionally, in the present study, *N. meningitidis* and *N. gonorrhoeae* were classified as the same species (BSC-sp7) due to their high genomic similarity (ANI values ≥ 95%) and patterns of gene flow. Defining bacterial species using arbitrary ANI thresholds or based on species name can lead to greatly overestimate amounts of introgression.

The variability of introgression levels are likely due to ecological and geographic factors. In some cases, such as *Haemophilus, Escherichia-Shigella*, *Campylobacter* and *Streptococcus*, we hypothesized that introgression may be the result of shared ecological niches. Cohabitating the same ecological niche such as a human host likely provides opportunities for frequent genetic exchange between related species. This idea is well supported by the patterns of introgression observed in the genus *Haemophilus* in which much of the species inhabit the same niche: the human respiratory tract and occasionally the genital area and gastrointestinal tract ^35–38^. Thus, niche overlap may play an important role in the prevalence of introgression between bacterial species. This is also the case of the genus *Campylobacter* with *C. coli* (BSC-sp28) and *C. jejuni* (BSC-sp3) colonizing the gut of humans and livestock, which could explain the high levels of introgression shared between these two species ^32,33,45^ (Supplementary Fig. 1k and Table 2). It has been hypothesized that *C. coli* and *C. jejuni* are genetically converging through introgression due to a recent ecological change ^54^. Under this model, introgression events represent a source of adaptive alleles transfers between *C. jejuni* and *C. coli* ^54^. The group *Escherichia-Shigella* includes species that colonize a broad host range including the human gut ^39–41^. The frequent introgression events observed between these enteric bacterial species may be driven by ecology. Conversely, introgression can potentially be a driver of ecological adaptation, which is potentially important for understanding genomic diversification of bacterial lineages such as the *Escherichia* species in the human gut. However, other factors are likely at play. Higher levels of introgression could be due to more permissive genetic machineries for recombination and to the presence and diversity of vectors capable of mediating gene flow. For instance, *Neisseria* and *Haemophilus* are capable of transformation and recognize species-specific DNA motifs ^55–60^. These motifs play crucial roles in homologous recombination by facilitating and controlling the process of exchange of genetic material between *Haemophilus* species. Similarly, *Vibrio* is capable of transformation and recognizes specific DNA motifs which are different from those described for *Haemophilus* ^61,62^. Natural transformation within the genus *Vibrio* may play a large part in the frequent homologous recombination occurring between the *Vibrio* species. *Streptococcus* species have been extensively studied for their ability to engage in natural transformation, and their genetic diversity and adaptivity could be the result of the frequent introgression that we observed between species within this genus ^63,64^.

DNA transfer is a mechanism that allows rapid adaptation and introgression likely contributes to this process. Our results suggest that introgression is likely adaptive by mediating the transfer of DNA sequences involved in certain functions such as carbohydrate and lipid metabolism. Although introgression events are restricted to the transfer of sequences between homologous genes, such transfers may be adaptive by allowing the fine tuning of gene function and expression to their environment. Our results revealed multiple instances of introgression events involving pathogenic bacteria such as *V. cholerae* and *V. alginolyticus*, *C. jejuni* and *C. coli* and several *Streptococcus* species. Introgression potentially plays a significant role in the evolution of bacterial pathogenicity. In addition, bacterial co-infections and super-infections involving related species may also offer opportunities for adaptation through increased virulence mediated by introgression.

As previously suggested ^12^, our results confirm that introgression is most common between highly related species. This may reflect the fact that homologous recombination requires high sequence relatedness to be efficiently processed ^31,65^. The fact that many related species present high levels of introgression can also be taken as evidence of ongoing bacterial speciation. According to some models, we expect that BSC-species could speciate through the progressive interruption of gene flow ^8,12,29,30^ Under this scenario, the observed patterns of introgression would correspond to decreasing gene flow due to ongoing speciation. However, it can be argued that introgression and ongoing speciation represent two sides of the same coin.

## Material and Methods

### Datasets

All analyzed genomes were downloaded from the GenBank database ftp.ncbi.nlm.nih.gov/genomes/. Lineages (*n*=51) were selected at the genus level based on the genomes available included in our previous study ^12^. All available genomes of each of those genera were downloaded from GenBank and included in the current study. This dataset included a total of 40,660 bacterial and archaeal genomes across 2,112 named species according to species designations on the NCBI website (Supplementary Data 1). Protein-coding genes of each genome were extracted based on the annotations. Note that all species with differing names at the time of this study were excluded (for example 118 species from the genus *Bacillus* used in the previous study ^12^ were re-named since, and those species were therefore excluded from this study). In addition, our overall dataset contained a single archaeal genus with 11 named species, and we therefore referred to our dataset as “bacteria” instead of “prokaryotes” to avoid generalization to all prokaryotes since our dataset includes a single archaeal genus. Finally, two genera, *Escherichia* and *Shigella* are highly related ^66^, sharing high sequence identity, and have been shown to be part of the same genus ^40,41,66^. All species from these two genera were therefore grouped into a single bacterial genus (named “*Escherichia-Shigella*”) resulting in a total of 50 genera in this study.

### Definition of core genomes and phylogenetic trees

For each genus, the core genome was built using *CoreCruncher* as performed previously ^67^ with *Usearch* Global v8.0 ^68^ and the stringent option. *CoreCruncher* was used because it can handle large datasets and because it includes a test to exclude potential paralogs and xenologs from the core genome (the inclusion of paralogs and xenologs into orthologous gene families becomes more likely as the number of genomes increases). Orthologs were defined with >70% protein sequence identity and >80% sequence length conservation and all other parameters were set to default. The core genome was defined as the set of single copy orthologs found in at least 85% of the genomes within each genus. Protein sequences of each core gene were then aligned using *Mafft* v7.407 ^69^ with default parameters. Protein alignments were then reverse-translated into their corresponding nucleotide sequences. Finally, the nucleotide alignments of all the core genes of each named species within the genus were concatenated into a single large alignment as previously described ^70^. Maximum likelihood phylogenomic trees were built from the concatenated alignment of the core genome for each genus using *FastTree* with the GTR model ^71^. Branch supports were evaluated by generating 1,000 bootstrap replicates using the same parameters. The trees were visualized with *FigTree* V1.4.4 (http://tree.bio.ed.ac.uk/software/figtree/).

### Species definition using the Average Nucleotide Identity (ANI) of the core genome

The core genome concatenates of each of the 50 genera were used to estimate the ANI of the core genome for all genome pairs. We used this method with a cutoff of 94% ANI as previously suggested ^34^, because the ANI of core genes is a slightly more stringent metric as core genes usually evolve slower than accessory genes ^12,34^. Pairwise ANIs of the core genomes were computed using the distmat tool of *EMBOSS v.6.6.0.0* ^72^, which calculates the pairwise nucleotide identities from the alignments ^34^. Then, single linkage clustering was performed as previously described ^34^: all genome pairs with an ANI of the core genome cutoff of 94% or higher were clustered together into a *de novo* species (ANI-species) (Table 1 and Supplementary Tables 1 and 2).

### Inference of BSC-species

We tested for the presence of gene flow between the core genomes of the pairs of ANI-species within each genus. Within each genus, all ANI-species with ≥ 15 genomes or more were the “reference species”. Species with fewer genomes available could not be used as reference species for this analysis. Then, we compared each reference species against one randomly selected genome of all other ANI-species from the genus which we named “candidate species”. For each comparison of a species against another one within the same genus, the core genome concatenate for the reference species + candidate species was re-built with *CoreCruncher* as described above to infer gene flow using the *ConSpeciFix* approach ^17^. Each core genome concatenate was used to compute a distance matrix using *RAxML* version 8.2.12 with the GTR+GAMMA model ^73^. From these distances, the ratio of homoplasic to non-homoplasic alleles (*h/m*) was computed for i) the reference species alone and ii) the reference species + the candidate species. Resampling analyses were conducted as previously described ^17^. From this step, graphs and statistics comparing *h/m* ratios between the genomes of each reference species with and without the candidate species were generated as previously described ^17^: The candidate species was inferred as a distinct BSC-species when a significant and substantial reduction of gene flow was detected based on *h/m* ratios (Wilcoxon test, *P*<0.0001). When no clear and significant reduction of gene flow was observed, the reference species and the candidate species were considered as putatively part of the same biological species and further tested for convergent mutations (see below).

### Convergent mutation test

Because our procedure is comparing various genomes, some comparisons can occasionally involve species with substantial genomic divergence. As genomes accumulate mutations during divergence, the frequency of convergent mutations increases as well, and this leads to the accumulation of homoplasic alleles that are the result of convergent mutations rather than gene flow. To account for this, we simulated genome sequence evolution with *CoreSimul* ^74^ for each dataset of reference species + candidate genome as described previously ^12^. The resulting simulated candidate genome sequence obtained for each pair of reference species + candidate species was evolved *in silico* with mutations but *without* gene flow and was used to estimate the ratio *h/m_0_* expected to result from convergent mutations alone against the ratios estimated for the reference species (Supplementary Table 3). The estimated values of *h/m_0_* were then compared to the real *h/m* values obtained between the candidate species and the reference species (*h/m_cand_*) as previously described ^12^. We considered cases where *h/m_0_* was similar to *h/m_cand_* as cases where the signal of gene flow is driven by convergent mutations rather than gene flow when *h/m_cand_* was not found significantly higher than *h/m_0_* (Wilcoxon test, *P*<0.0001). In such cases, the reference and candidate species were considered as distinct species. This step was conducted for each pair of reference species and candidate species within each genus (22,894 comparisons).

### Inference of introgression events

Introgression events were inferred based on the combination of phylogenetic signal and sequence identity. We used a phylogenetic approach to infer candidate introgression events where we compared individual gene trees to core genome trees. Introgression events were defined as sequences of core genes exchanged by homologous recombination between distinct species (the analysis was conducted separately for ANI-species and BSC-species). To infer introgressed core genes, unrooted trees were built for each core gene using *RAxML* ^73^ with a GTR + GAMMA model for each genus. For each genus, the topology of each unrooted gene tree was compared to the topology of the unrooted genome tree. Each ANI-species and each BSC-species were tested for monophyly from the unrooted core genome tree. Non-monophyletic species (*n*=50), as inferred from the core genome phylogeny, were ignored for this analysis. A putative introgression event of a core gene was inferred when the sequences of the ANI-species (or the BSC-species) were not found to be monophyletic in the unrooted gene tree (see Fig. 1). When these introgression events were inferred, it was sometimes possible to unambiguously determine which species pairs engaged in introgression with one another (i.e., when the sequence of a single species was nested within the subtree of a different monophyletic species). To further ascertain the inference of introgression events in our dataset, we restricted our analysis to pairs of BSC-species where both BSC-species have at least 15 genomes because we cannot infer BSC-species with high confidence when less than 15 genomes are available (Supplementary Table 2). We only inferred introgression events between species pairs when at least one of the species pairs was found not engaging in gene flow with the other BSC-species. Moreover, although Incomplete Lineage Sorting (ILS) has not been described in bacteria, it has been shown that this mechanism can be confounded for introgression in animals and plants ^75^. However, introgression is expected to leave a distinct signature from ILS: because introgressed genes involve the transfer of a sequence from one species to another, introgression events will theoretically lead to higher sequence identity of the introgressed sequence shared by the two species. In contrast, higher sequence identity is not expected under ILS. Therefore, the candidate introgression events inferred based on phylogenetic incongruencies were confirmed as introgression events when the tested gene displayed substantially higher sequence identity than the overall divergence of the core genome between the two species. For each pair of BSC-species, we calculated the highest identity score for each core gene between the pairs of species, the average identity score of the sequences of that gene between the pairs of BSC-species and the standard deviation (SD). Similar trends of introgression were inferred between BSC-species for each genus when using a threshold of 2 SD, 3 SD, and 4 SD. The threshold of 2 SD was then selected to infer introgression events between BSC-species within each genus (Supplementary Fig. 4).

### Functional annotation of core genomes into COG categories

To determine whether introgressed genes were biased toward certain functions, the core genes of each genus were annotated and grouped into functional categories by comparison to the COG (Cluster of Orthologous Genes) database using eggNOG-mapper v2.1.4 ^49^. The ratio of genes assigned to each COG category against the total number of genes in the core genome and the introgressed genes in each genus was analyzed, respectively. Within each genus, only the pairs of BSC-species sharing at least 10 introgressed genes were selected to avoid biased data. A Wilcoxon test with Bonferroni *P*-value adjustment was conducted to compare the distribution of genes involved in each COG category to the genes in all other COG categories between the introgressed genes and the entire core genome across genera and among all the genera. All other BSC-species pairs (and genera composed only of BSC-species pairs sharing < 10 introgressed genes) were excluded from the comparison.

## Acknowledgments

This study was supported by the National Science Foundation NSF grant DEB-1831730 (LMB) and the National Institutes of Health grant R01GM132137 (LMB).

## Author contributions

A.D. and LM.B. contributed to the conception and design of the project, performed analyses, and wrote the manuscript.

## Data availability

Genomes used in this study are listed in Supplementary Data 1 and are freely available on GenBank at https://www.ncbi.nlm.nih.gov/genome/. All the core genomes, the genome trees and the gene trees generated in the present study are available on Kaggle at https://www.kaggle.com/datasets/awadiop/diop-nature-ecology-evolution-2024-1 https://www.kaggle.com/datasets/awadiop/diop-nature-ecology-evolution-2024-2 https://www.kaggle.com/datasets/awadiop/diop-nature-ecology-evolution-2024-3

## Supplementary information

**Supplementary Fig. 1. Core genome phylogenetic analysis of the 50 genera.** Maximum likelihood phylogenomic trees were built from the concatenated alignment of the core genome for each genus. Percentage bootstrap support values below 99% are shown for each node. The scale of the branch length of the trees (bottom) are expressed in nucleotide substitutions per site. Each colored box with a number represents an ANI-species. The named species associated with each ANI-species are indicated on the right. The trees were annotated when multiple ANI-species where reclassified into a single BSC-species. **a**, *Acinetobacter*. **b**, *Actinomyces*. **c**, *Aeromonas*. **d**, *Alteromonas*. **e**, *Bacillus*. **f**, *Bacteroides*. **g**, *Bartonella*. **h**, *Bifidobacterium*. **i**, *Bordetella*. **j**, *Burkholderia*. **k**, *Campylobacter*. **l**, *Chlamydia*. **m**,*Citrobacter*. **n**, *Clostridium*. **o**, *Corynebacterium*. **p**, *Cronobacter*. **q**, *Escherichia-Shigella*. **r**, *Elizabethkingia*. **s**, *Enterobacter*. **t**, *Enterococcus*. **u**, *Flavobacterium*. **v**, *Fusobacterium*. **w**, *Gilliamella*. **x**, *Haemophilus*. **y**, *Helicobacter*. **z**, *Klebsiella*. aa, *Lactobacillus*. **ab**, *Lactococcus*. **ac**, *Leptospira*. **ad**, *Methanosarcinia*. **ae**, *Mycobacterium*. **af**, *Mycobacteroides*. **ag**, *Neisseria*. **ah**, *Paenibacillus*. **ai**, *pantoea*. **aj**, *Photobacterium*. **ak**, *Prevotella*. **al**, Pseudomonas. **am**, Rhizobium. **an**, *Rhodococcus*. **ao**, *Salmonella*. **ap**, *Serratia*. **aq**, *Sinhorizobium*. **ar**, *Staphylococcus*. **as**, Stenotrophomonas. **at**, *Streptococcus*. **au**, *Vibrio*. **av**, *Xanthomonas*. **aw**, *Yersinia*.

**Supplementary Fig. 2. Levels of introgression across ANI-species and genera. a**, Percentage of introgressed core genes across pairs of ANI-species within each genus. **b**, Relationship between levels of introgression and core genome sequence identity across pairs of ANI-species.

**Supplementary Fig. 3. Distribution of percentage of introgressed core genes across genera.**

**Supplementary Fig. 4. Inference of introgressed genes based on sequence identity.** For each pair of BSC-species, each dot represents a core gene. The graphs compare the average sequence identity of the core gene between the two BSC-species to the highest sequence identity score of the gene between both BSC-species. The black line represents *y=x*. Red dotted lines represent the threshold of 2 standard variation (2 SD). Red dots represent the core genes inferred as introgressed based on tree incongruency. Green dots represent non-introgressed core genes whose tree was congruent with the tree of the core genome.

**Supplementary Fig. 5. Functional annotations of introgressed genes across genera. a**, Classification of core genes into COG categories among all genera. **b**, Classification of introgressed genes into COG categories across genera. Stacked bars represent the distribution of genes in each COG category for each genus. See the legend key for the description of each COG category.

**Supplementary Fig. 6. Functions of introgressed genes relative to the core genome composition for each genus.** *P* values of COG categories were calculated with a Wilcoxon test and were adjusted for multiple testing using a Bonferroni correction. See the legend key for the description of each COG category.

**Supplementary Table 1.** Percentage of introgressed core genes across species and genera.

**Supplementary Table 2.** Introgression and Average Nucleotide Identity among ANI-species and BSC-species.

**Supplementary Table 3.** Metrics inferred for the reclassification of the ANI-species into BSC-species.

**Supplementary Data 1:** List of the genomes and strains of the 50 genera included in this study.

